# Sassy2: Batch Searching of Short DNA Patterns

**DOI:** 10.64898/2026.03.10.710811

**Authors:** Rick Beeloo, Ragnar Groot Koerkamp

**Affiliations:** Department of Theoretical Biology and Bioinformatics, Utrecht University, Karlsruhe, Germany; Department of Computer Science, Karlsruhe Institute of Technology, Karlsruhe, Germany

**Keywords:** Approximate string matching, pattern matching, fuzzy string searching, semi-global alignment, edit distance, bitpacking, SIMD, DNA

## Abstract

**Motivation:** Searching short DNA patterns such as barcodes, primers, or CRISPR spacers within sequencing reads or genomes is a fundamental task in bioinformatics. These problems are instances of multiple approximate string matching (MASM) [Baeza-Yates and Navarro, 1997], which requires locating all occurrences with up to *k* errors of multiple patterns of length *m* in a text of length *n*. Classical approaches based on seeding with exact matches become inefficient for short patterns (*m ≤* 64 bp) as *k* increases, producing either many spurious hits or missing true matches. Our previous work, `Sassy1`, showed that careful hardware optimization drastically accelerates single-pattern searches in long texts by distributing chunks of the text across SIMD lanes.

**Methods:** `Sassy2` distributes multiple *patterns* across SIMD lanes to maximize parallelism when searching batches of short patterns. When *k* is small, often only a short substring of the pattern of length *O*(*k*) is needed to reject a possible match. Thus, `Sassy2` first examines short suffixes of the patterns (e.g., the last 16 bp of 32 bp patterns), allowing more (but smaller) parallel SIMD lanes. Only positions passing this suffix filter undergo full pattern verification.

**Results:** On synthetic data, `Sassy2` achieves 10–50*×* speedups over `Sassy1` for short texts (*n ≤* 200 bp) and 2–4*×* for large texts (*n ≥* 1 Mbp). On real-world tasks with 16 threads, `Sassy2` reaches over 100 Gbp/s text throughput per guide when searching 312 gRNAs across the human genome and 116 Gbp/s throughput when demultiplexing Nanopore reads with 96 barcodes. In both cases, `Sassy2` outperforms `Sassy1` by 2–5*×* and Edlib by 20–45*×*.

**Availability:** `Sassy2` is implemented in Rust and available at github.com/RagnarGrootKoerkamp/sassy.

## 1. Introduction

### Short patterns in biology

Many tasks in bioinformatics require searching multiple short DNA patterns (e.g. 20 − 40 bp) within larger sequences. Examples include demultiplexing Nanopore barcodes [Beeloo et al., 2025], finding CRISPR off-target sites [Wang et al., 2024], primer sequences [Ye et al., 2012], and others [Nicolae and Rajasekaran, 2015].

### From exact to approximate matching

Early sequence analysis relied on exact matching algorithms such as Knuth–Morris–Pratt [Knuth et al., 1977] and Boyer–Moore [Boyer and Moore, 1977], which respectively achieve *O*(*n*) and *O*(*n/m*) time complexity to search a pattern of length *m* in a text of length *n*. However, biological variations and sequencing errors require algorithms that tolerate mismatches, insertions, and deletions [Smith et al., 1981b]. While classical dynamic programming methods provide such error-tolerant alignments, they require *O*(*nm*) time, which is impractical for large-scale applications [Needleman and Wunsch, 1970, Smith et al., 1981a].

### Myers’ bit-vector algorithm

A major breakthrough in approximate string matching (ASM) came in 1999 with Myers’s bit-vector algorithm [Myers, 1999], which encodes dynamic programming (DP) matrix columns into machine words and updates them using bitwise operations, computing edit distance in *O*(*n*?*m/w*?) time, where *w* is the word size (typically 64). The algorithm uses what we will call *pattern tiling*: at each step, *w* pattern characters are compared simultaneously against a single text character. For short patterns (*m* = *O*(*w*)), this achieves *O*(*n*) time in practice.

### Optimizations for short patterns and low error thresholds

Pattern tiling becomes inefficient when *m* ≪ *w*, because many bits in the word remain unused. To mitigate this, Fredriksson [2003] proposed *text tiling*, which compares a single pattern character against *w* text characters in parallel, as also implemented in `Sassy1` [Beeloo and Groot Koerkamp, 2025].

Returning to *pattern tiling*, another idea for scenarios with many short patterns is *pattern tiling with packing*: several short patterns are concatenated in a machine word, allowing multiple patterns to be processed simultaneously. If *r* patterns satisfy 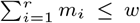, they can be searched in parallel [Hyyrö et al., 2005]. For example, when *m* ≈ *w/*2 ≈ 32 bp, packing two patterns yields an approximate 2× speedup, with larger gains as *m* ≪ *w*. In practice, however, many biological patterns are ≈ 20 bp long, and thus only a few patterns can be packed into a single word.

A key observation is that the entire pattern rarely needs to be compared in order to reject a random alignment at a small edit-distance threshold *k*. This idea is commonly formalized by *L*(*k*), the expected number of dynamic-programming (DP) rows that must be computed before a random alignment is rejected. Building on this, Hyyrö et al. [2005] proposed packing only the first *L*(*k*) (approximately 2*k*) characters of each pattern and continuing with the full Myers’ algorithm only when the packed prefix indicates a promising match. Since *L*(*k*) ≪ *m* for small *k*, this strategy increases packing density while keeping verification costs low when true matches are rare. For a broader overview of related optimizations, see Wu and Manber [1992] and Hyyrö et al. [2005].

### From intra-word to SIMD lane parallelism

In the 1980s, and 1990s, many optimizations focussed on the *word-size*. However, with modern hardware more bits are available. Specifically, SIMD (Single-Instruction-Multiple-Data) instructions allow parallelism across multiple independent words, eliminating the need for complex intra-word packing. A SIMD register of bit-width *W* consists of *L* lanes of width *w* (*W* = *L* · *w*). For example, `AVX2` provides 256-bit registers supporting lane configurations (*w, L*) ∈ {(8, 32), (16, 16), (32, 8), (64, 4)}. The AVX2 instruction set is widely available on modern x86 CPUs, while newer architectures may support `AVX-512`, which extends the register width to 512 bits. In `Sassy1` we used text-tiling where a single text is split into *L* equal-sized chunks that are distributed over SIMD lanes, each holding their own Myers state. When *n* = *O*(*W*), this becomes inefficient as bits remain unused and parts of the matrix are computed twice at overlapping chunk boundaries. In practice, `Sassy1` only reaches its maximal throughput for *n* ≥ 8000. Instead, in `Sassy2` we use SIMD-optimized pattern-tiling where batches of *L patterns* (*m* ≤ 64) are searched against the same text.

Many biological patterns naturally fit these word sizes, making SIMD a practical alternative to intra-word parallelism.

### Our contributions

We present a practical SIMD implementation of multi-pattern ASM using Myers’ bit-packing that is substantially faster than previous approaches. Specifically:

1. **Multi-pattern SIMD implementation:** Multiple short equal-length patterns are packed into SIMD lanes with the pattern encodings optimized for SIMD loading (Appendix A.1). Patterns are matched in parallel, with parallelism determined by the SIMD width and allowed number of errors.
2. **Suffix filter:** We first search short (length just above 2*k*) suffixes of all patterns using smaller word sizes (and thus, more SIMD lanes), and only verify full-pattern matches when the suffix matches with cost ≤ *k*. This is simpler and computationally cheaper than the early-break check in `Sassy1` and substantially improves throughput for typical applications.

## 2. Methods

In this work we extend `Sassy1` with optimized batch pattern searching, to which we refer as `Sassy2`. We adopt the notation of Beeloo and Groot Koerkamp [2025]. A *pattern P* has length *m* and a *text T* has length *n*, both defined over alphabet Σ. The *Multiple Approximate String Matching* (MASM) problem is: given a set of patterns P = {*P*_1_, …, *P*_*r*_}, an error threshold *k*, and text *T*, find all end positions *j* in *T* such that *d*(*P, T* [*i* … *j*]) ≤ *k* for some *P* ∈ P and some 0 ≤ *i* ≤ *j*, where *d*(·, ·) denotes unit-cost Levenshtein (edit) distance. We use *L*(*k*) to denote the expected number of DP rows to be computed to reject a random pattern against a random text.

### 2.1. SIMD-accelerated bit-parallelism

Our implementation builds on Myers’ bit-parallel algorithm [Myers, 1999], which encodes DP matrix columns into machine words of width *w*, resulting in per text character updates in *O*(*m/w*) word-sized operations. We extend this with SIMD registers of bit-width *W*, partitioned into *L* independent lanes of width *w* (so *W* = *L*·*w*). Each lane maintains an independent Myers bit-vector state.

#### SIMD text-tiling (long texts)

In `Sassy1` we used *text-tiling* [Beeloo and Groot Koerkamp, 2025] for long texts (*n* ≫ *W*), partitioning the text into *L* chunks and processing them in parallel against a single pattern. This approach allows early rejection since rows of the DP matrix are computed one by one: often only the first ≈ *L*(*k*) rows (corresponding to pattern characters) need to be computed to reject a mismatch [Ukkonen, 1985, Beeloo and Groot Koerkamp, 2025], reducing complexity from *O*(*m* · ⌈*n/w*⌉) to *O*(*L*(*k*) · ⌈*n/w*⌉) when *k* is small.

#### SIMD pattern-tiling (many short patterns)

Text-tiling becomes inefficient for short texts (*n* = *O*(*W*)) or multiple patterns: partitioning short texts leaves SIMD registers underutilized, while multiple patterns require repeated text rescanning and encoding. We therefore use *pattern-tiling*, encoding all patterns once and distributing them across *L* SIMD lanes (of width *w*) to compare each text character against all patterns simultaneously (Figure 1).

**Fig. 1.**
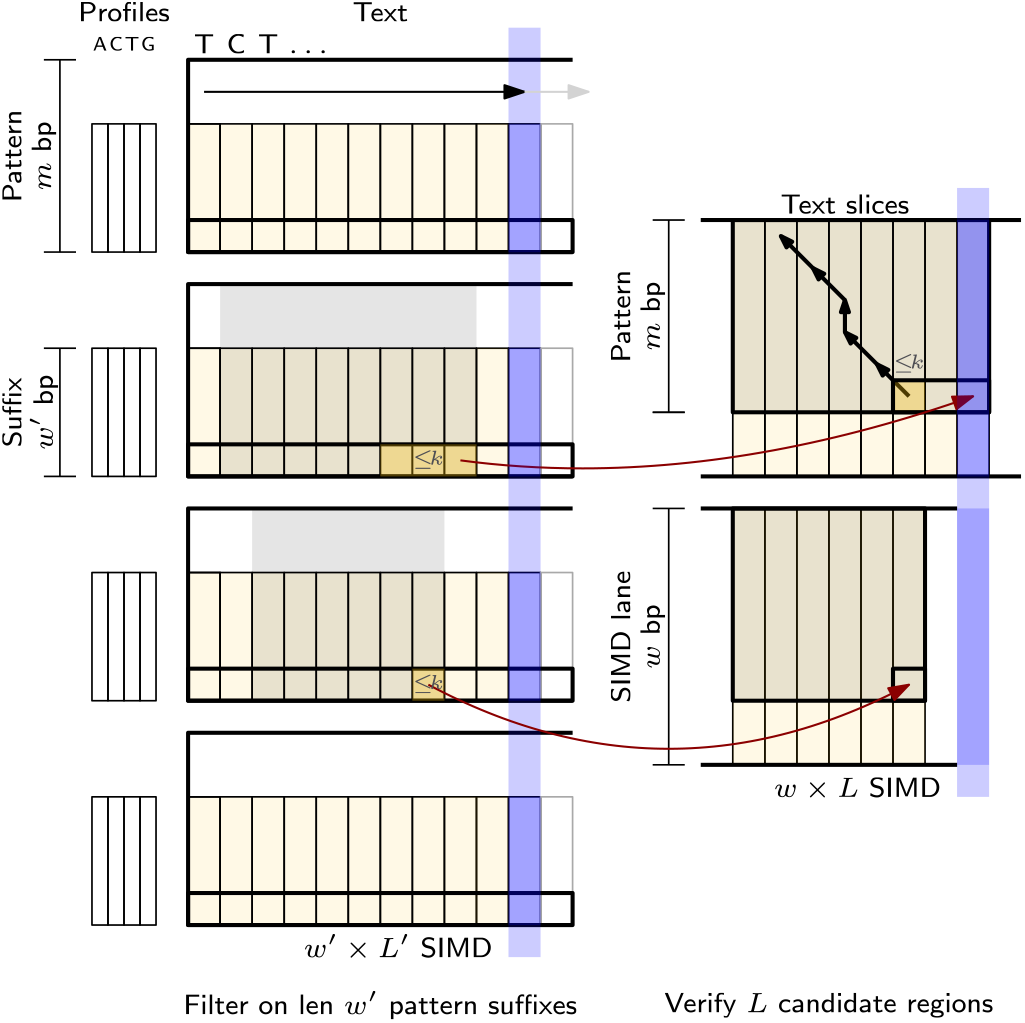
SIMD pattern tiling overview. First, the full patterns and their suffixes of length *w*′ bp are encoded using a *profile* (e.g DNA or IUPAC). Then, the text characters are processed one by one (left to right) and each character is compared against *L*′ pattern suffixes of length *w*′ bp using a *w*′ *× L*′ SIMD tiling (blue vertical bar, Section 2.2, shown for 4 *×* 4). The end positions of suffixes that match with *≤ k* errors are highlighted in yellow. When such positions are adjacent they form a single range *R*. We then verify whether the full pattern matches with *≤ k* errors at those end positions. The slices corresponding to each suffix range (2 ranges here), each of size |*R*| + *m* + *k −* 1, are then in parallel verified against the full pattern using a *w × L* tiling (here 2 *×* 8). The local minima (single opaque cell) with value *≤ k* are selected for traceback (small black arrows) to report the matching text locations and CIGAR string.

Each lane maintains independent Myers bit-vector state, enabling *L* parallel pattern searches. This scans the text once rather than *r* times (once per pattern). However, unlike in text-tiling, the full pattern is compared at every position, so early rejection based on the first *L*(*k*) characters is not directly possible (Figure 1). We address this through suffix filtering.

### 2.2. Suffix filtering

To recover early rejection while preserving pattern-tiling, we use a two-stage approach: a fast suffix filter running at reduced lane width *w*′ *< w*, followed by full pattern verification at width *w*. This relates to Hyyrö et al. [2005]’s prefix filtering with *L*(*k*) but differs in: (1) using suffixes (rather than prefixes) for direct endpoint traceback (discussed later), (2) empirically-tuned *w*′ ∈ {8, 16, 32} rather than theoretical *L*(*k*), though we expect *L*(*k*) ≈ *w*′, and (3) processing across *L* parallel lanes rather than packing multiple patterns within a word.

#### Stage 1: Suffix filter

We first search only pattern suffixes using reduced lane width *w*′ ∈ {8, 16, 32}, which increases the number of lanes (*L*′ = *W/w*′ *> L*) and thus parallelism (Figure 1, left). The suffix filter maintains the same error threshold *k*. We select *w*′ based on *k* (Table 1): *w*′ = 8 for exact matching (*k* = 0), *w*′ = 16 for few errors (0 *< k <* 4), and *w*′ = 32 for moderate errors (4 ≤ *k <* 8). When *w*′ equals the original lane width (e.g., for very short patterns or large *k*), no filtering is applied.

**Table 1.** Timings (in milliseconds) of different suffix filters (*w*′) for a given pattern length *m*, using word size *w*, with edit distance cut-off *k*. For each configuration, 96 random DNA patterns are queried against a fixed text of length *n* = 1000, and runtimes are averaged over 1000 iterations. When *w*′ = *w*, the filter is skipped. The fastest time for each (*m, k*) pair is shown in bold.

| $m$ | $k$ | $w$ | $w' = 8$ | $w' = 16$ | $w' = 32$ | $w' = 64$ |
| --- | --- | --- | --- | --- | --- | --- |
| 16 | 0 | 16 | **0.019** | 0.028 | – | – |
| 16 | 1 | 16 | 0.089 | **0.028** | – | – |
| 16 | 2 | 16 | 1.546 | **0.030** | – | – |
| 16 | 3 | 16 | 11.930 | **0.031** | – | – |
| 16 | 4 | 16 | 42.584 | **0.099** | – | – |
| 23 | 0 | 32 | **0.018** | 0.030 | 0.057 | – |
| 23 | 1 | 32 | 0.137 | **0.030** | 0.058 | – |
| 23 | 2 | 32 | 2.077 | **0.031** | 0.058 | – |
| 23 | 3 | 32 | 17.117 | **0.045** | 0.058 | – |
| 23 | 4 | 32 | 58.702 | 0.081 | **0.076** | – |
| 23 | 8 | 32 | 108.322 | 51.594 | **0.556** | – |
| 32 | 0 | 32 | **0.021** | 0.031 | 0.061 | – |
| 32 | 1 | 32 | 0.173 | **0.031** | 0.064 | – |
| 32 | 2 | 32 | 3.081 | **0.030** | 0.062 | – |
| 32 | 3 | 32 | 21.846 | **0.037** | 0.061 | – |
| 32 | 4 | 32 | 76.123 | 0.104 | **0.061** | – |
| 32 | 8 | 32 | 139.036 | 65.412 | **0.061** | – |
| 64 | 0 | 64 | **0.019** | 0.029 | 0.059 | 0.116 |
| 64 | 1 | 64 | 0.448 | **0.050** | 0.069 | 0.149 |
| 64 | 2 | 64 | 5.691 | **0.030** | 0.060 | 0.119 |
| 64 | 3 | 64 | 40.444 | **0.036** | 0.058 | 0.120 |
| 64 | 4 | 64 | 138.812 | 0.210 | **0.065** | 0.115 |
| 64 | 8 | 64 | 259.417 | 131.087 | **0.060** | 0.121 |

#### Stage 2: Full pattern verification

Each suffix match with cost ≤ *k* at text position *i* is a candidate that requires full pattern verification. Any full pattern match ending at *i* must lie within the text slice *s* = *T* [*i* − *m* − *k* … *i*], since up to *k* deletions can shift the alignment start by at most *k* positions. We compute the full Myers DP matrix over *P* × *s*, storing column-wise deltas for traceback. Using SIMD, we can re-compute the DP matrix for up to *L* end-points in parallel (fig. 1, right). Then, all end-points for which the full pattern matches with ≤ *k* errors are optionally filtered depending on the match reporting mode (section 2.3), after which we perform a traceback to obtain the CIGAR string, cost, and text interval.

#### Batch tracing

When multiple suffix endpoints of cost ≤ *k* cluster together, we group them into contiguous ranges for joint verification (highlighted cells in Figure 1). This is expected as edit distance is “smooth”: horizontally adjacent DP states differ by at most 1 in edit distance, so nearby endpoints with cost ≤ *k* naturally form clusters. A given range *R* starts at *R*_*s*_ till *R*_*e*_ (exclusive) with length |*R*|, and corresponds to the text slice *T* [*R*_*s*_ … *R*_*e*_]. We then compute the matrix *P* × *T* [*R*_*s*_ − *m* − *k* + 1 … *R*_*e*_] and store the deltas for each processed text character. This amortizes DP construction: instead of computing independent matrices totaling |*R*| · (*m* + *k*) columns, we compute only |*s*| = |*R*| + *m* + *k* − 1 columns.

### 2.3. Match reporting

We support two match reporting modes [Beeloo and Groot Koerkamp, 2025]: **(i)** report all end positions in the text where an alignment of cost ≤ *k* ends, or **(ii)** report only rightmost local minima ≤ *k*. Both modes are implemented for batch searching. Like in `Sassy1`, we also support *overhang costs* allowing reduced cost for unaligned regions at the start or end of the text, see Beeloo and Groot Koerkamp [2025] for more details.

### 2.4. Time complexity

The suffix-filter processes *r* suffixes of length *w*′ in batches of *L*′ = *W/w*′ lanes, where *W* is the SIMD register width. Each batch executes a constant number of SIMD instructions per text character, giving a total cost of Θ(*n*?*r/L*′?). The *C* candidates that pass the suffix filter are verified with worst-case complexity *O*((*m* + *k*)?*C/L*?), where *k* is the number of allowed errors and *L* = *W/w* is the verification parallelism. In practice, the filter is highly selective (with only few false positives) and the first term dominates unless there are many matches.

## 3. Results

We compare `Sassy2` to `Sassy1`, based on AVX2, and `Edlib`, a widely used edit-distance search library without SIMD vectorization [Šošić and Šikić, 2017]. All methods are evaluated in their *library* form, excluding I/O effects and reflecting purely algorithmic performance. Unlike the others, `Edlib` performs semi-global alignment and returns only minimum-cost alignments. This allows `Edlib` to dynamically tighten the threshold *k* during the search, improving runtime, but not reporting matches whose cost is non-minimal but still ≤ *k*. All experiments are conducted on an Ubuntu machine with an XEON GOLD 6530 CPU (32 cores, 64 threads, running at 4 GHz, with AVX-512 support). Patterns are only searched in the forward direction^1^.

### 3.1. Synthetic data

We compare the tools on random synthetic data using a single thread while varying the text length *n* and the number of patterns *r* (Figure 2). Unless noted otherwise, results are for AVX-512.

**Fig. 2.**
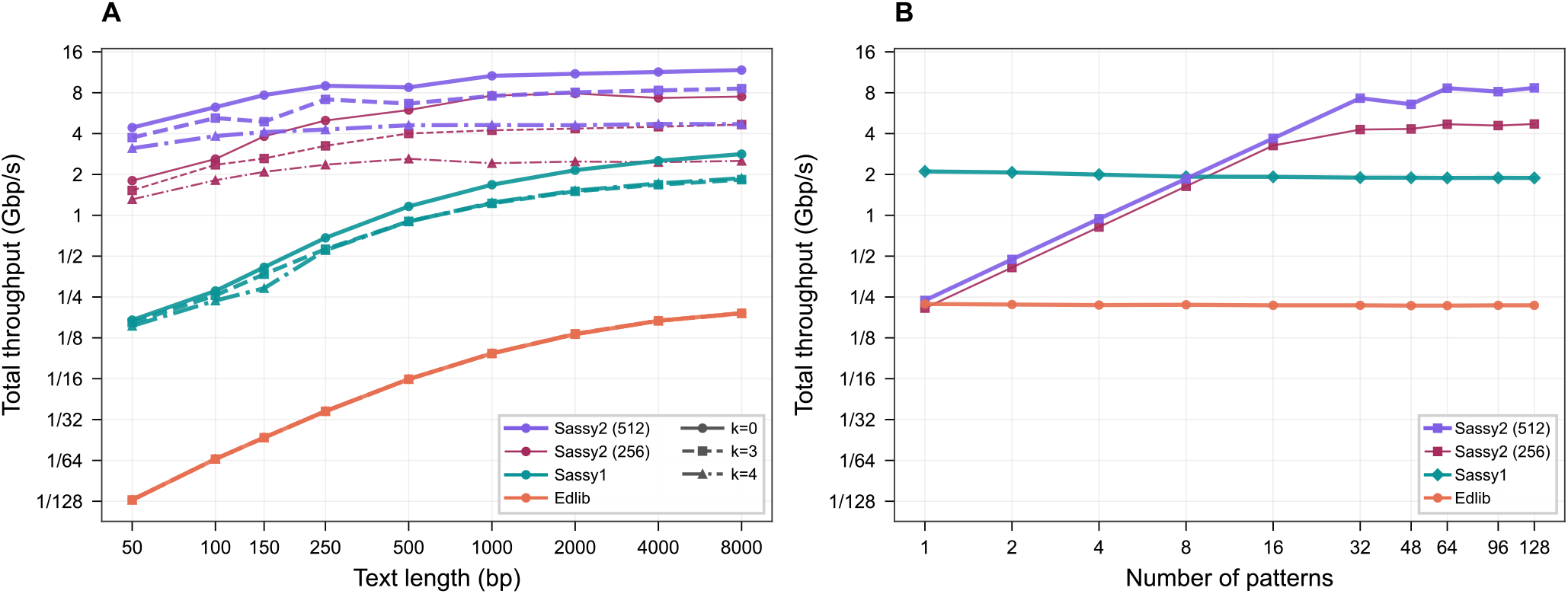
Performance comparison of approximate string matching algorithms. **(A)** Throughput (Gbp of text per second per pattern; log_2_ scale) for `Sassy2` with AVX-512 (512-bit SIMD; purple) and AVX2 (256-bit SIMD; dark pink), along with `Sassy1` (pattern tiling; teal) and Edlib (coral). Experiments search *r* = 128 patterns of length *m* = 23 in random text of varying length *n ∈ {*50, …, 10000*}*, with edit distance thresholds *k ∈ {*0, 3, 4*}*. For `Sassy2`, batch throughput is multiplied by the number of patterns *r* to report total throughput per pattern. **(B)** Scaling with the number of patterns *r* (log_2_ scale) for *m* = 23, *n* = 100,000, *k* = 3 with throughput as shown for (A).

#### Scaling with text length

We search *r* = 128 patterns of length *m* = 23 bp with up to *k* ∈ {0, 3, 4} errors and vary *n* from 50 bp to 8 kbp, corresponding to short texts, typical Illumina reads (150 bp), and long Nanopore reads (8 kbp), see Figure 2A. For *k* = 3, `Sassy2` achieves 3.74, 4.90, and 8.58 Gbp/s for these lengths respectively. This corresponds to speedups of 23×, 13×, and 4.6× over `Sassy1`, and 467×, 213×, and 45× over `Edlib`. Unlike `Sassy1`, which is slow for short texts, `Sassy2` is nearly as fast for short texts as for long texts. Thus, `Sassy2` provides significant gains when searching, for example, short reads.

#### Scaling with the number of patterns

We search an *n* = 100 kbp text with 1 to 128 patterns (Figure 2B) of length *m* = 23 bp with *k* = 3. Total throughput increases linearly from 0.24 to 7.29 Gbp/s as *r* grows from 1 to 32 and more SIMD lanes are used, and then plateaus. At *r* = 32, this corresponds to a 3.85× speedup over `Sassy1` and 33.6× speedup over `Edlib`.

#### Effect of SIMD width

For *m* = 23 and *k* = 3, `Sassy2` uses a suffix filter of length *w*′ = 16 (Table 1). With AVX2, this allows up to 16 suffix comparisons per step, and with AVX-512 up to 32. Accordingly, Fig. 2B shows near-linear throughput scaling with the number of patterns until the SIMD lanes are fully filled with suffixes, after which scaling flattens. AVX2 achieves proportionally lower peak throughput (e.g. 2.62 vs. 4.90 Gbp/s at *m* = 150; Figure 2A), consistent with its smaller register width.

#### Overall performance

`Sassy2` outperforms `Edlib` by nearly three orders of magnitude on small texts and achieves substantial gains over `Sassy1` once SIMD lanes are fully utilized.

### 3.2. Real-world applications

To evaluate `Sassy2` on real-world workloads, we performed (i) CRISPR off-target searching in the human genome and (ii) barcode detection in Nanopore reads. These experiments use 16 threads and patterns are searched in the forward strand only.

#### CRISPR off-target searching

Minimizing off-target effects is a central step in CRISPR editing. Guide RNAs (gRNAs) must therefore be searched genome-wide for matches with up to *k* errors. We search 312 guide RNAs (23 bp) from the CRISPRoffT database [Wang et al., 2024] against the complete CHM13v2.0 human genome (3.12 Gbp) allowing up to *k* = 3 errors, a common off-target threshold [Modrzejewski et al., 2020]. With the suffix filter, `Sassy2` reaches amortized 105.9 Gbp/s *per pattern*, corresponding to an amortized wall-clock time of 30 ms per guide. In comparison, `Sassy1` reaches 28.6 Gbp/s (109 ms per guide, 3.7× slower) and Edlib 3.0 Gbp/s (1051 ms per guide, 35.7× slower).

#### Barcode searching in Nanopore reads

A common demultiplexing task for Nanopore reads is to find all barcodes with up to *k* errors [Beeloo et al., 2025, Cheng et al., 2024]. We search 96 barcodes of 24 bp from the rapid barcoding kit (`SQK-RBK114.96`) in Nanopore reads from *Photobacterium acropomis* (`SRR26425221`) [Gould and Henderson, 2023]. This dataset is 334 Mbp, with 37 bp to 97 kbp reads (3.7 kbp avg.).

For *k* = 3, with *w*′ = 16 suffix filter, `Sassy2` achieves amortized 116.8 Gbp/s per barcode with a *total* wall-clock time of 0.27 s to search the 96 patterns in 334 Mbp of reads. In comparison, `Sassy1` reaches 25.5 Gbp/s (1.26 s, 4.6× slower) and `Edlib` 2.57 Gbp/s (12.5 s, 45× slower). At *k* = 4, without suffix filter, `Sassy2` reaches 58.7 Gbp/s per barcode (0.55 s), still 2.3× faster than `Sassy1` (25.4 Gbp/s, 1.26 s) and 23× faster than `Edlib` (2.56 Gbp/s, 12.5 s).

#### Overall performance

Across real-world genomics tasks, `Sassy2` searches at over 100 Gbp/s per pattern with the suffix filter enabled, and remains at least 2× faster than `Sassy1` and `Edlib` without it, enabling rapid, high-sensitivity searches across whole genomes or long Nanopore read sets.

## 4. Discussion

We introduced `Sassy2`, a SIMD-accelerated tool for batch searching of short DNA patterns that combines Myers’ bit-vector algorithm with multi-lane SIMD parallelism in the *pattern direction* and a suffix filter.

On synthetic data, `Sassy2` achieves up to 467× speedup over `Edlib` and up to 23× over `Sassy1` for short texts (*n* = 50 bp), and 44× and 4.5× speedups, respectively, for longer texts (*n* = 10 kbp). Throughput scales near-linearly with the number of patterns until SIMD lanes are saturated (e.g., 7.3 Gbp/s at *r* = 32, *n* = 10 000 bp, AVX-512).

On real-world genomics data, `Sassy2` reaches 105.9 Gbp/s per pattern with 16 threads to search 312 gRNAs in the human genome and 116.8 Gbp/s per pattern for 96 Nanopore barcodes. Even without the suffix filter, it remains at least 2× faster than `Sassy1` and 20× faster than `Edlib`, demonstrating high-throughput searches across genomes and long reads.

### Limitations

`Sassy2` currently requires patterns of equal length, whereas `Sassy1` supports variable-length searches. This limitation currently has to be worked around by processing patterns of different lengths separately. Combining the suffix filter of `Sassy2` with text tiling is a promising direction for future work.

### Conclusion

`Sassy2` extends our previous SIMD-optimized approach for long texts to efficient batch searching of short equal-length patterns, providing a practical, high-throughput solution for a wide range of genomics search tasks and overall demonstrating that early ideas can be adopted to modern hardware.

## A. Algorithm implementation details

### A.1. Pattern-tiling encoding

We begin with the standard Myers construction for a *single* pattern. Consider the IUPAC alphabet of 16 symbols encoded into 4 bits (0 ≤ *c <* 16), with each bit indicating whether the symbol matches A/C/G/T. For a pattern *P* of length *m* and each symbol *c* ∈ {0, …, 15}, we precompute a bitmask

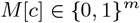

where bit *j* is set if symbol *c* matches *P* [*j*]. During scanning, for each text position *i*, we encode the character (*c* = encode(*T* [*i*])) and use *M* [*c*] in the Myers update. Here encode returns the 4 bit encoding of the ASCII character *T* [*i*].

For multiple patterns 𝒫 = {*P*_1_, …, *P*_*r*_}, the natural extension is to construct masks

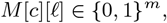

for every character *c* and pattern index *ℓ* ∈ {0, …, *r* − 1}, where bit *j* is set if *c* matches position *j* of *P*_*ℓ*_. If these masks are stored pattern-by-pattern, then for each text character *T* [*i*] we have to iterate over all *ℓ* and load *M* [*c*][*ℓ*] from separate memory regions. This is inefficient as we cannot load all the “equality masks” for a given character *c* into the *L* different SIMD lanes directly. To enable SIMD pattern-tiling, we therefore *transpose* the layout. Let the SIMD register width be *W* = *L*·*w*, consisting of *L* independent *w*-bit lanes. We group patterns into SIMD blocks of size *L* and define

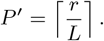

For each character *c*, pattern *ℓ* is stored in block

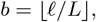

and within the block in lane

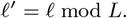

Thus the corresponding SIMD vector is

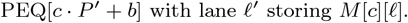

This ensures that each SIMD vector contains the equality masks for *L* different patterns at the same character *c*, packed by lane. Then during scanning, for each text position *i* we compute *c* = encode(*T* [*i*]) once and obtain a pointer to the contiguous region PEQ[*c* · *P* ′ … (*c* + 1) · *P* ′). The inner loop then streams linearly over the *P* ′ SIMD blocks, loading one equality vector per block and applying the *standard* Myers update independently in each lane.

### A.2. Suffix filter benchmarking

Table 1 shows benchmarks determining the optimal word-size *w*′ for the suffix filter.

## Footnotes

1 `Sassy2` and `Sassy1` both support reverse-complement searching, but ASM is not invariant: `Sassy1` searches patterns in the reverse-complement of the text, while `Sassy2` searches the reverse-complement of the patterns in the forward text, yielding potentially different matches [Beeloo and Groot Koerkamp, 2025].

